# Contrasting reproductive strategies despite convergent traits for two dryland river-floodplain species

**DOI:** 10.1101/2022.05.12.491621

**Authors:** Will Higgisson, Linda Broadhurst, Foyez Shams, Bernd Gruber, Fiona Dyer

**Affiliations:** Centre for Applied Water Science, University of Canberra, University Drive, Bruce, Canberra, ACT 2617, Australia; Centre for Australian National Biodiversity Research, CSIRO National Research Collections Australia, GPO Box 1700, Canberra, ACT 2601, Australia; Centre for Conservation Ecology and Genetics, University of Canberra, University Drive, Bruce, Canberra, ACT 2617, Australia

**Keywords:** Propagule biology, Aquatic plants, Semi-arid and arid, Floodplain, Dispersal, Reproduction

## Abstract

Aquatic plants share a range of convergent reproductive strategies, such as the ability to reproduce both sexually and asexually through vegetative growth. In dryland river systems, floodplain inundation is infrequent and irregular, and wetlands consist of discrete and unstable habitat patches. In these systems life-history strategies such as long-distance dispersal, seed longevity, self-fertilisation, and reproduction from vegetative propagules are important strategies which allow plants to persist. Using two aquatic plants *Marsilea drummondii* and *Eleocharis acuta*, we investigated the proportions of sexual and asexual reproduction and self-fertilisation employing next generation sequencing approaches and used this information to understand population genetic structure in a large inland floodplain, in western New South Wales (NSW) Australia. Asexual vegetative reproduction and self-fertilisation was more common in *M. drummondii*, but both species used sexual reproduction as the main mode of reproduction. This resulted in highly differentiated genetic structure between wetlands and similar genetic structure within wetlands. The similarity in genetic structure was influenced by the wetland in the two species highlighting the influence of the floodplain landscape and hydrology in structuring population genetic structure. The high levels of genetic variation among wetlands and low variation within wetlands suggests that dispersal and pollination occur within close proximity and that gene flow is restricted. This suggests a reliance on locally sourced (persistent) seed, rather than asexual (clonal) reproduction or recolonisation via dispersal, for population maintenance in plants in dryland rivers. This highlights the importance of floodplain inundation to promote seed germination, establishment and reproduction in dryland regions.

## Introduction

Aquatic plants are an extremely heterogeneous group of species sharing a range of convergent reproductive and other life history traits which allow them to live and thrive in aquatic environments (Philbrick and Les 1996; Li 2014). One of the distinguishing reproductive strategies shared by most aquatic plants is asexual reproduction which includes the production of seed without fertilization (agamospermy) and vegetative reproduction (Philbrick and Les 1996). Although vegetative reproduction is common in aquatic plants, most species retain the ability for both sexual and asexual reproduction (e.g. *Phragmites australis* (Liu *et al*. 2021), *Decodon verticillatus* (Dorken and Eckert 2001) and *Cladium jamaicense* (Ivey and Richards 2001)). The challenges of successful fertilisation in aquatic environments have also seen the evolution of some unique strategies such as hydrophily where pollen is transported under water (Les 1988), and pollination and seed set within closed flowers (eg. in Callitriche, Philbrick and Les 1996).

Sexual reproduction increases genetic diversity, while asexual reproduction maintains genetic uniformity of well-adapted gene complexes (Les 1988; Niklas and Cobb 2017). The higher genetic variability in sexual propagators facilitates adaption for long-term persistence whereas asexual propagators promote short-term adaption which can be beneficial for rapid establishment and expansion (Li 2014). Li (2014) proposes that asexual reproduction in aquatic plants is a strategy that ensures population maintenance, while sexual reproduction is responsible for population restoration from extreme events at evolutionary timescales. If this is the case, we may expect that aquatic plants will have highly differentiated populations with little among-population gene flow and extensive within-population clonality.

Asexual reproduction provides an alternative reproductive strategy when sexual recruitment is challenging or limited. Barrett (1980) observed that in *Eichhornia crassipes* (water hyacinth) sub-optimal seed production was related to in-efficient pollinator service and seedlings were only recorded at sites with saturated soil and shallow water conditions. In the aquatic macrophyte *Phragmites australis* (common reed) seed production was found to be highly variable between sites and years and was related to rainfall and temperature (McKee and Richards 1996). Interspecific variation in sexual and asexual reproduction was observed in the aquatic plant *Decodon verticillatus* (swamp loosestrife) where asexual reproduction became more dominant toward the geographical range limits of the species. This was related to reduced sexual performance related to genetic (such as reduced pollen tube growth and ovule penetration) and ecological factors (primarily changes in temperature) (Dorken and Eckert 2001). For this species, traits involved in sexual reproduction in the northern population contributed little to fitness and, as such, are likely to have been disabled through genetic mutation (Eckert *et al*. 1999).

In vegetative (asexual) reproduction, there are two levels of population structure which can provide meaningful data on plant demography; the ‘ramet’ which is the unit of clonal growth (i.e. a single individual plant produced by clonal propagation), and the ‘genet’ is the colony of all the genetically indistinct ramets (i.e. there can be many ramets of a genet) (Harper 1977). Sampling clonal species can be challenging and structured methodologies to maximise the sampling of genets are required. This is to ensure that an accurate representation of the number of discrete individuals and their genotypes within a population are obtained. The organs of clonal propagation (for example tillers, rhizomes or bulbs) influence the spatial arrangement and size of ramets and the amount of outcrossing which may occur between ramets (Barrett 2015). The size of a clone has been found to influence sexual reproduction through pollen deposition, with larger clones found to be increasingly saturated with pollen arriving from the same clone in *Carex platyphylla* (broad leaf sedge) (Handel 1985).

A range of factors are likely to influence population genetic structure in aquatic plants such as hydrology (Junk *et al*. 1989) and the river and floodplain topography as well as plant morphology, pollination systems (Eckert *et al*. 2016), modes and rates of extinction, dispersion and colonisation (Wade 1978), and sexual and asexual propagule dispersal (Boedeltje *et al*. 2004) and establishment (Van der Valk 1981). While rivers facilitate the movement of individuals and genes (Junk *et al*. 1989), high levels of genetic differentiation among populations and low within-population variation is often observed in aquatic plants, and maybe related to widespread asexual (clonal) reproduction (Santamaría 2002). Significant genetic differentiation between populations was observed in *Cladium jamaicense* (swamp sawgrass) with genetic structure reflecting patterns from colonisation maintained through long-lived clones (Ivey and Richards 2001). Similarly, Piquot *et al*. (1996) observed genetically discrete populations consisting of a single or few clones in *Sparganium erectum* (branching bur-reed).

One reason for the high genetic differentiation often observed among populations is due to long-distance dispersal (>100m) being rare in plants (Le Corre *et al*. 1997; Cain *et al*. 2000). Despite this rarity, long-distance dispersal does occur and is essential for colonising new suitable habitats (Cain *et al*. 2000). It is of particular importance for unstable local populations which may be more prone to localised extinction events (Hanski 1998). The flow regime of rivers in dryland regions is highly variable, where floodplain inundation is infrequent and irregular and reliant on rainfall during particularly wet years (Walker *et al*. 1995). This means that floodplain wetlands may be discrete and possibly unstable habitat patches (Hanski 1998) with aquatic plants in these habitat patches naturally going through localised extinction and colonisation events. Here long-term population survival relies on long-distance dispersal (Cain *et al*. 2000; Ozinga *et al*. 2004) and ability of seed to remain dormant and viable between flooding events. If floodplain habitats in dryland regions are meta-populations, extinction rates and modes (and rate) of recolonisation (either from the entire metapopulation (migrant-pool) or from a single source population (propagule-pool)) will influence population genetic structure (Wade 1978; Pannell and Charlesworth 1999). Repeated extinction and population turn-over in metapopulations for example, reduces within-population diversity and increases among population diversity, especially in the absence of migration between extant populations (Pannell and Charlesworth 1999).

The genetic distinctiveness of individuals has long been a challenge in plant demography especially for species that are clonal where plant census are unsuitable (Harper 1977). Recent advances in genomics have made the identification of ramets and their genets more achievable allowing the extent of clonality in aquatic plants such as *Laminaria rodriguezii* (kelp) (Reynes *et al*. 2020) and *Chara* spp. (Schaible *et al*. 2009) to be determined. Understanding the population genetic structure of aquatic plants provides information including the importance of sexual and asexual reproduction to guide and predict the consequences of land or water management actions on these species.

The Murray-Darling Basin covers an area of 1 million km^2^ in the southeast of Australia and is an important region of Australia for agricultural production as well as supporting around 2.3 million people and a range of environmental assets (MDBA 2012). Many rivers that make up the Murray-Darling Basin occur in semi-arid and arid regions and are characterised by low and extreme variability in flows (Walker *et al*. 1995). These systems have undergone pronounced changes over the last century through river regulation (Kingsford 2000; Higgisson *et al*. 2020).

Two aquatic/semi-aquatic plants which reproduce sexually and asexually via vegetative growth that are common on the floodplains of the Murray-Darling Basin are *Marsilea drummondii* A. Braun (common nardoo) and *Eleocharis acuta* R.Br. (common spike-rush). On the floodplains of the Murray-Darling Basin, *M. drummondii* and *E. acuta* are often the dominant groundcover during and after flooding events, forming monospecific stands or clumps (Pers Obs. W Higgisson and F Dyer). Using these two species, we investigated the extent and spatial arrangement of 1) asexual vegetative propagules, 2) self-fertilisation, and 3) plants in a parent-offspring relationship across a large inland floodplain in western New south Wales (NSW), Australia using next generation sequencing approaches. Using this information, we also explored the population genetic structure of both species. Understanding the population genetic structure of these plants provides important information on mating patterns, resource allocation and the role of hydrology in the maintenance and establishment of these and other plants on floodplains in dryland regions.

## Methods

### Study area

This study was carried out in three wetlands on the floodplain of the lower Lachlan River, NSW, Australia (Figure 1), where both species occur. The lower Lachlan River has a semi-arid climate which experiences a mean annual rainfall of 319 mm (at Oxley, NSW (station number: 049055) Bureau of Meteorology 2021a). The temperature ranges from a mean monthly Summer maximum of 35.1°C in January to a mean monthly Winter maximum of 15.8°C in July (at Hay (Airport), NSW, (station number: 075019) Bureau of Meteorology 2021b).

**Figure 1.**
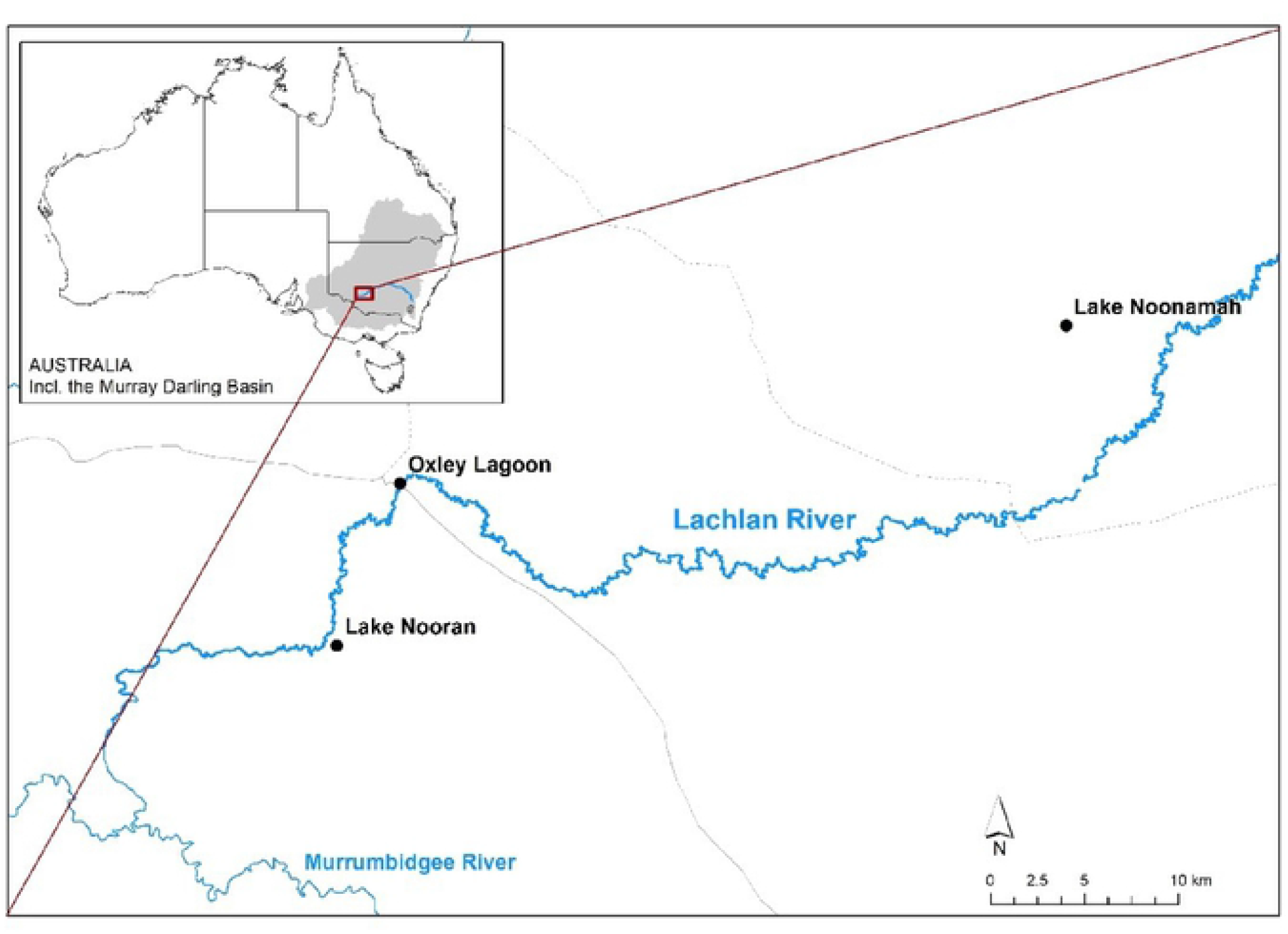
*The floodplain wetland sites used in this study where* Eleocharis acuta *and* Marsilea drummondii *samples were collected from within 2-3 patches in each wetland. The inset shows the location of the Lachlan River (in blue) and the Murray-Darling Basin (shaded light grey) in Australia.*

### Study species

#### Marsilea drummondii

*Marsilea drummondii* (Marsileaceae) is an aquatic/semi-aquatic perennial fern widely distributed across inland mainland Australia (PlantNET 2021a). It is a submerged plant with leaves floating on the water surface and occurs at the edge of lakes, swamps and floodplains. The fronds are clover-like, growing at the end of stalks which vary in length from 2-30 cm.

Marsileaceae are aquatic and semi-aquatic heterosporous perennial, rooted water ferns forming rhizomatous spreading clumps (Jones 1998). Ferns such as this alternate between haploid and diploid phases with the diploid phase being the most visible. *Marsilea* produce a modified seed-like spore-bearing leaf known as a sporocarp which has a thick sclerenchymatous wall that protects the enclosed sori against damage such as by insect attack and dryness (Nagalingum *et al*. 2006). When stimulated by water the sporocarp produces sporangia which gives rise to haploid male microspores and female megaspores with internal fertilization giving rise to the next diploid generation. *Marsilea drummondii* sporocarps are 4-9 mm long, on stalks 8-35 mm long (Cunningham *et al*. 1981).

Reproduction in *Marsilea* is highly dependent on the flooding regime and wetting and drying cycles, with the entire reproductive biology dependent on water for the release of spores from the sporocarp, dispersal of spores following fertilization and sinking of young developing embryos (Schneider and Pryer 2002). Sporocarp fruiting in *Marsilea* (including in *M. drummondii*) occurs primarily on mud as a habitat dries (Johnson 1986). Viable sporocarps of several *Marsilea* species have been reported to germinate from herbarium material up to 100 years old (Lellinger 1985) highlighting the ability of this genera to remain viable for long periods. *Marsilea drummondii* has germinated in soil seedbank studies, indicating their ability to remain viable and contribute to the soil seedbank (Higgisson *et al*. 2021 unpublished).

The sporocarp can be transported by adhesion, such as to the external surfaces of birds and can be ingested (Johnson 1986). The sporocarps of *M. mucronata* have been shown to pass intact through the digestive tract of waterbirds (Malone and Proctor 1965). The size and weight of sporocarps (sporocarps in *M. drummondii* range from 4-9 mm long) is predicted to limit wind dispersal (Jones 1998). Hydrochory is likely to be an important dispersal mechanism for sporocarps, during flooding events, however sporocarps start to open within a few hours of contact with water (Schneider and Pryer 2002). Once the sporocarp opens the spores are short-lived and must germinate quickly and are highly susceptible to drying, therefore spores are unlikely to be units of long-distance dispersal and dormancy (Schneider and Pryer 2002).

#### Eleocharis acuta

*Eleocharis acuta* (Cyperaceae) is a rhizomatous perennial plant which grows to 90 cm high (Cunningham *et al*. 1981; PlantNET 2021b). *Eleocharis acuta* occurs along the eastern and south western mainland of Australia and across Tasmania as well as New Zealand, New Guinea, and Norfolk Island (PlantNET 2021b). It grows in moist situations along streams, roadsides, floodplains and around dams.

*Eleocharis acuta* seeds can float on water (50% of seeds floated for > 30 days), and the species is likely to disperse via hydrochory; seed are dormant although the mechanism involved is unknown, and require extended contact with water to germinate (> 40 days) (Higgisson and Dyer 2021). Intact *E. acuta* seeds were recorded in waterbird fecal samples by Green *et al*. (2008) suggesting that waterbirds are seed dispersers. All members of the Cyperaceae are wind pollinated (Walters 1949). While the timing of spikelet maturity is unknown for *E. acuta*, the spikelet of *E. laeviglumis* from southern Brazil is dichogamous and protogynous with reproductive organs maturing at different times (Demeda *et al*. 2018) which reduces self-pollination and promotes out-crossing.

### Study design and sampling strategy

Leaf samples of *M. drummondii* and *E. acuta* were collected from Lake Nooran, Oxley Lagoon and Lake Noonamah. Lake Nooran is an open temporary wetland a few hundred metres from the main channel of the Lachlan River in a complex of wetlands where the Lachlan River terminates, Oxley Lagoon is in close proximity (<50 m) to the Lachlan River up-stream of Lake Nooran, while Lake Noonamah is on an ephemeral channel off the main stem of the Lachlan River (Figure 1). Within each wetland, a total of 13 leaf samples were collected from within 2 or 3 patches (Figure 2) that were approximately 100 m apart. Sampling at each patch consisted of 13 points starting from a central point at which a single leaf was collected. From the central point the nearest leaf was collected from the point 0.5 m North, East, South, and West, then 1.5 m from the central point, and finally 2.5 m from the central point in the same pattern (Figure 2). A total of two patches were sampled within Lake Nooran and Oxley Lagoon and three patches at Lake Noonamah for each species. This resulted in a total of 91 samples for each species.

**Figure 2.**
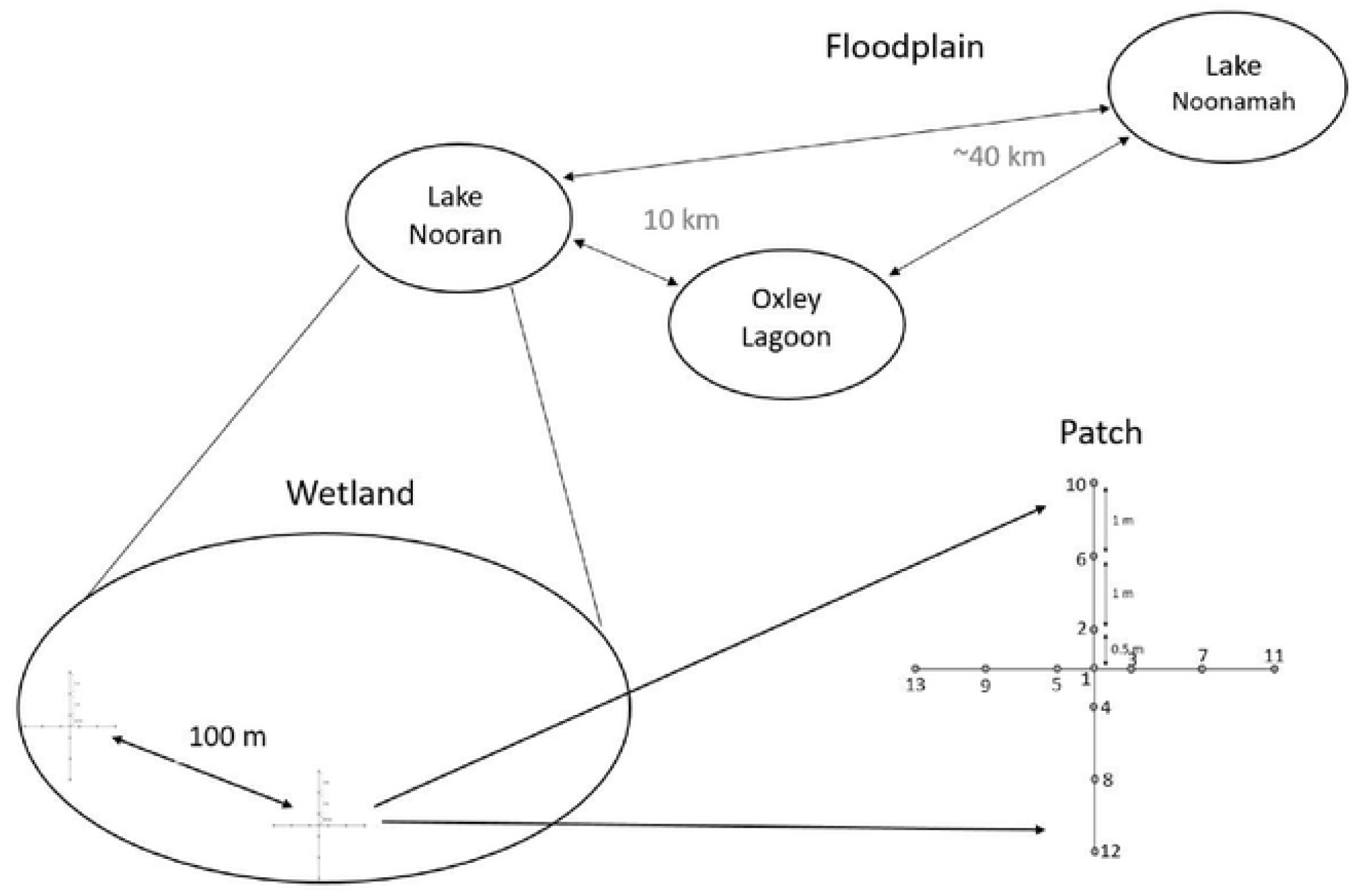
*Conceptual diagram showing the three wetlands used in this study, how patches were arranged within each wetland and how samples were collected within each patch.*

### Genotyping

DNA extraction was performed using protocols developed by Diversity Array Technology Pty Ltd (DArTseq™). Tissue samples (25 mg) were incubated overnight at 56° C with lysis buffer and proteinase K. Lysed tissue was washed using washing buffer to remove impurities such as protein and polysaccharides and stored in elution buffer. Genotyping was undertaken on samples from each species using a genome wide profiling approach using restriction enzymes to produce DNA fragments, which were pooled (for each species) and sequenced using next-generation platforms by Diversity Arrays Technology (http://www.diversityarrays.com). DArTseq™ represents a combination of a DArT complexity reduction method and next generation sequencing platforms (Kilian *et al*. 2012; Cruz *et al*. 2013). Therefore, DArTseq™ represents an implementation of sequencing complexity reduced representations (Altshuler *et al*. 2000) and more recent applications of this concept on the next generation sequencing platforms (Baird *et al*. 2008; Elshire *et al*. 2011). A detailed description of the DArTseq™ methodology can be found in Killian *et al*. (2012). DarTseq genotyping produced 45,940 single nucleotide polymorphisms (SNPs) for *M. drummondii* and 150,801 SNPs for *E. acuta*.

### Data analysis

The SNP data supplied by DarTseq and associated metadata were read and filtered using the software package dartR, Version 1.9.1 (Gruber *et al*. 2018). As this research was interested in determining the relatedness of samples and their spatial arrangement within and among wetlands, we initially investigated pairwise frequencies in consistent loci. We determined three levels of pairwise relatedness being 1) genetically identical pairs through asexual clonal reproduction; 2) pairs in a parent-offspring relationship via self-fertilisation; and 3) pairs in a parent-offspring relationship via out-crossing (Table 1). Under these three pairwise relatedness scenarios the true number of inconsistent loci in each scenario should be zero. Since it is possible that PCR error might result in small differences between identical samples, we filtered the data based on a read depth ≤12 and ≥50 for *Marsilea drummondii* and a read depth ≤15 and ≥50 for *Eleocharis acuta* (considering the much larger starting dataset) which removed or significantly reduced the occurrence of SNP errors in the datasets. We also filtered the data on average repeatability of alleles at a locus of 99.8%, a call rate threshold of 0.95; and all secondary SNPs where then removed. This filtering left 1,241 SNPs for *M. drummondii* and 6,618 SNPs for *E. acuta* which were used to determine the three levels of pairwise relatedness.

**Table 1.**
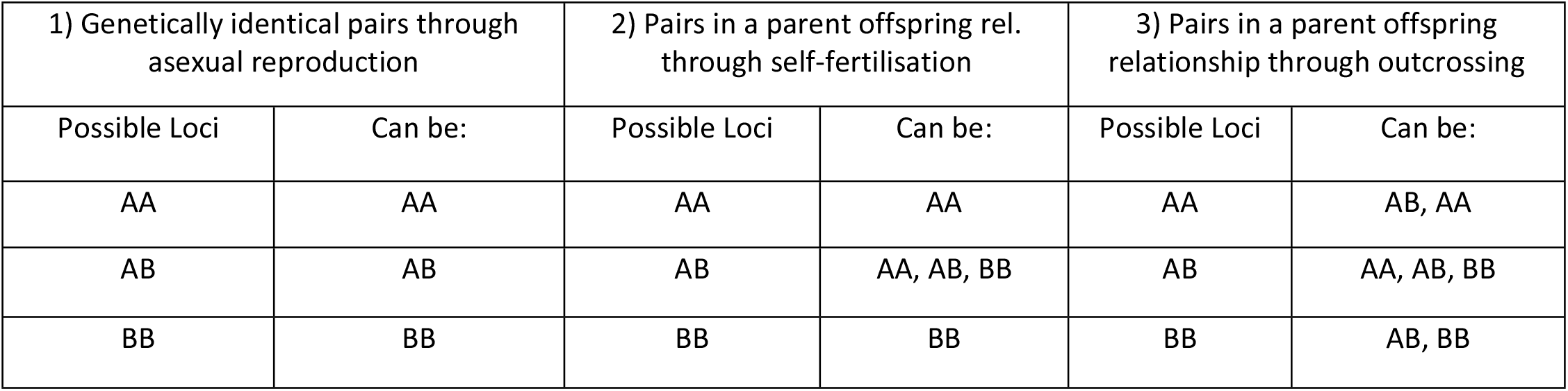
pairwise relatedness scenarios used as part of this study and the possible options at a given loci.

| 1) Genetically identical pairs through asexual reproduction |  | 2) Pairs in a parent offspring rel. through self-fertilisation |  | 3) Pairs in a parent offspring relationship through outcrossing |  |
| --- | --- | --- | --- | --- | --- |
| Possible Loci | Can be: | Possible Loci | Can be: | Possible Loci | Can be: |
| AA | AA | AA | AA | AA | AB, AA |
| AB | AB | AB | AA, AB, BB | AB | AA, AB, BB |
| BB | BB | BB | BB | BB | AB, BB |

While we filtered the SNP data stringently to reduce sequencing error, it may still have been possible that PCR error might result in small differences between pairs within each of the relatedness scenarios. Therefore, we have provided the results from the relatedness analysis for each species as Appendix a for *E. acuta* and Appendix b for *M. drummondii* showing the number of inconsistent loci and the total number of loci used in the analysis between each pair of samples in each of the three relatedness scenarios. We report only the pairs with no (0) loci differences in the results but highlight that those pairs with a single or few loci differences may also be in the relatedness scenarios.

Following the relatedness analysis, a single representative of each genotype (i.e., the genetically identical pairs determined in scenario 1) from each patch was retained (i.e., one of each clonal group per patch) leaving 84 *M. drummondii* and 90 *E. acuta* genotypes for further analysis. These datasets were then re-filtered as described above to produce a dataset of 1,242 SNPs for *M. drummondii* and 6,623 SNPs for *E. acuta*.

Observed (*H*_o_) and expected (*H*_e_) heterozygosity was calculated for each species across all samples, within each wetland and within each patch using the function gl.report.heterozygosity from the package dartR (Gruber *et al*. 2018). An inbreeding coefficient (*F*_IS_) was calculated for each patch for each species, using the function boot.ppfis in the package Hierfstat, Version 0.5.7 (Goudet 2005), which uses the formula *F*_IS_ = 1-(*H*_o_/*H*_e_), with 1000 bootstraps.

Principle coordinates analysis (PCoA) was undertaken on all genotypes for each species in dartR, Version 1.9.1 (Gruber *et al*. 2018). Genetic structure was estimated using the program Structure, version 2.3.4 (Pritchard *et al*. 2000). The number of populations tested was assumed to be ‘k’ = from 1 to 10. Using an admixture model the analysis was run 10 times for each ‘K’ value with 10000 burnin length and 10000 MCMC replication after burnin. The optimal values of ‘K’ (number of panmictic clusters) were identified for each species following Evanno’s method (Evanno *et al*. 2005) using Structure Harvester (Earl 2012). Overall genetic differentiation was estimated using Analysis of Molecular Variance (AMOVA) undertaken using the function poppr.amova in the package poppr, Version 2.9.2 (Kamvar *et al*. 2014) using the ade4 implementation of AMOVA. The level of genetic differentiation among patches and among wetlands was assessed by calculating the *F*_ST_ values using the package GenAlEx, Version 6.5 (Peakall and Smouse 2006).

## Results

### Clonality

The 91 samples for each species collected from two patches in Lake Nooran and Oxley Lagoon and three patches in Lake Noonamah (13 samples per patch) yielded 83 genotypes for *M. drummondii* and 90 for *E. acuta* (Table 2). Seven *M. drummondii* samples from within Lake Nooran patch 1 were genetically identical with samples distributed over an area of approximately 2 m^2^ (see appendix c for spatial arrangement of samples). There were also three genetically identical *M. drummondii* samples from Lake Noonamah, two from patch 1 (samples 1 and 12 which were 2.5 metres apart) and the third from patch 2 (Appendix c). In *Eleocharis acuta*, there was one pair of samples that were genetically identical in Lake Noonamah patch 2 (50 cm apart). No other patches contained clones in either species.

**Table 2.** *Number of genotypes within each wetland and patch for* Marsilea drummondii *(MD) and* Eleocharis acuta *(EA). The numbers in brackets are the number of samples collected.*

|  |  | (n) genotypes |  |  |  |
| --- | --- | --- | --- | --- | --- |
| Wetland | Patch | MD<br>Wetland | Patch | EA<br>Wetland | Patch |
| Lake Nooran | 1 | 21 (26) | 7 (13) | 26 (26) | 13 (13) |
|  | 2 |  | 13 (13) |  | 13 (13) |
| Noonamah<br>Wetland | 1 | 37 (39) | 12 (13) | 38 (39) | 13 (13) |
|  | 2 |  | 13 (13) |  | 12 (13) |
|  | 3 |  | 13 (13) |  | 13 (13) |
| Oxley Lagoon | 1 | 26 (26) | 13 (13) | 26 (26) | 13 (13) |
|  | 2 |  | 13 (13) |  | 13 (13) |
| All-Lachlan |  | 83 (91) |  | 90 (91) |  |

### Parent-offspring relationship through self-fertilisation

For *M. drummondii*, we only detected pairwise samples in a parent-offspring relationship via self-fertilisation in Lake Noonamah (Appendix a) where one sample in patch 1 was related to three other patch 1 samples, with 4 m distance between these samples (Appendix c). A sample in Lake Noonamah patch 2 was observed to be in a parent-offspring relationship via self-fertilisation with two samples from patches 1 and 2. There were three Lake Noonamah patch 3 samples in a parent-offspring relationship via self-fertilisation, which were also in a parent-offspring relationship via self-fertilisation with a further four samples from the other two Lake Noonamah patches (3 samples in one patch and 1 in the other).

In *Eleocharis acuta*, there was only one pair of samples in a parent-offspring relationship via self-fertilisation in Lake Noonamah (100 cm apart) (Appendix d).

### Parent-offspring relationship through outcrossing

In *M. drummondii* a total of 422 (or 10% of the total) pairs were in a parent-offspring relationship through outcrossing (Appendix a). The majority of these pairs occurred within and between patches within a wetland with slightly more than half (52%) of these occurring between samples within a patch, and ∽47% occurring between samples in different patches within the same wetland. Three samples in Oxley Lagoon patch 2 were in a parent offspring relationship through outcrossing with three samples in Lake Nooran. In *E. acuta*, only 34 pairs of samples were in a parent-offspring relationship. These pairs only occurred within patch and not between patches or wetlands.

### Other sample pairs with few loci differences

Results from pairwise relatedness analysis based on the three relatedness scenarios for *M. drummondii* (Appendix a) and *E. acuta* (Appendix b) also highlighted that there were some genotype pairs in both species with only a single or few loci differences in the three relatedness scenarios. In *M. drummondii* there were 12 pairs that had a single loci difference in the clonal relatedness scenario and a further 28 pairs that had a single loci difference in the parent-offspring by self-fertilisation scenario (of 1238 loci). In *E. acuta* there were three pairs that had between 2 and 5 inconsistent loci (out of 6588 loci) in the clonality scenario while the next most similar pair had 41 loci differences (see appendix b). These pairs could be genetically identical through clonal reproduction or in a parent-offspring relationship through self-fertilisation.

### Genetic diversity

Observed and expected heterozygosity varied between the two species. *Marsilea drummondii* had a mean observed heterozygosity (*H*_*o*_) across all genotypes of 0.105 compared with *E. acuta* which had a mean *H*_*o*_ of 0.165 (Table 3). Despite this difference, similar patterns in *H*_*o*_ and *H*_*e*_ were observed between the two species across the different wetlands and patches. For example, *H*_*o*_ and *H*_*e*_ was lowest for both species at Lake Noonamah compared to the same measures at Lake Nooran and Oxley Lagoon. Further, at the patch level, *H*_*o*_ and *H*_*e*_ were the lowest at Lake Noonamah patch 3 compared with the other patches. *H*_*e*_ was the greatest in both species in Oxley Lagoon and within Oxley Lagoon patch 1 at the patch level. The highest *H*_*o*_ was observed in Oxley Lagoon in *M. drummondii* and Lake Nooran in *E. acuta* (Table 3). The inbreeding coefficient (*F*_IS_) in *M. drummondii* was greater than 0 at Lake Noonamah patches 1 and 2, and slightly greater than 0 at Oxley Lagoon patch 2 which indicated inbreeding has occurred at these patches. In *E. acuta, F*_IS_ was less than 0 in all patches, demonstrating excess heterozygosity as compared to expected in all patches, and widespread occurrence of outcrossing (Table 3).

**Table 3.**
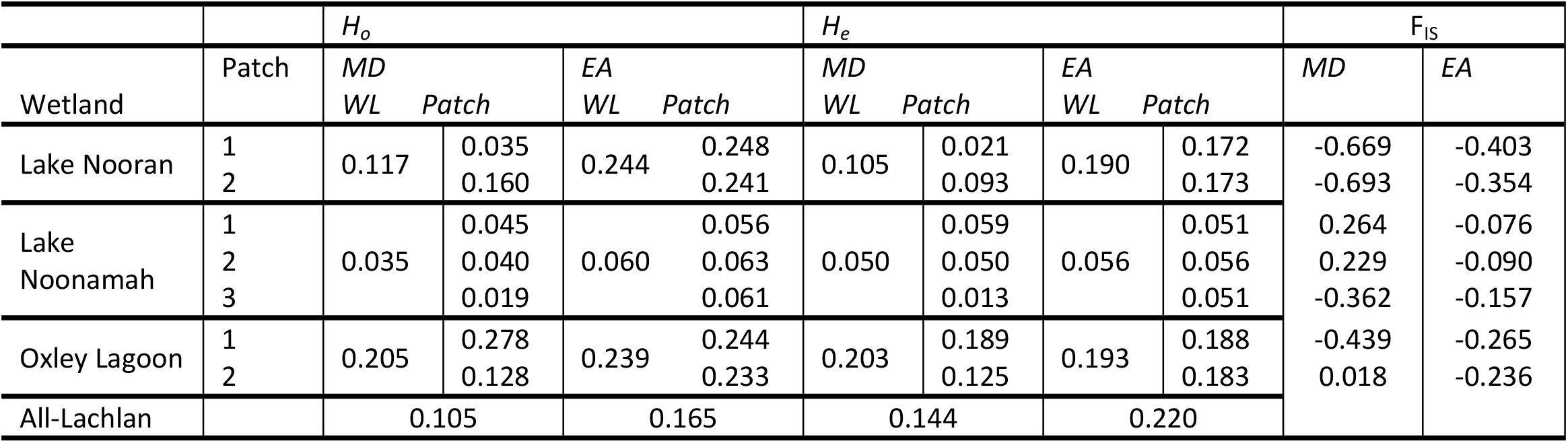
*Observed (H*_*o*_*) and expected (H*_*e*_) *genetic diversity within wetlands, patches and overall for* Marsilea drummondii *(MD) and* Eleocharis acuta *(EA). WL = wetland. F*_*IS*_ *= (median) Inbreeding coefficient.*

### Population genetic structure

The first two PCoA axes accounted for 57% of the total variation in *M. drummondii* with similar variation along both axes (31.9% vs. 25.2%). Most samples were clustered together within the wetland patch from which these were collected (Figure 3a) with several discrete clusters present. At the site level Lake Noonamah plants were differentiated from all others and fell into two clusters with the exception of one sample that fell near a small group of Lake Nooran plants. The remaining Lake Nooran plants grouped with some of the Oxley Lagoon plants with the remainder of clustering some distance away. At the patch level most plants were grouped together with some exceptions such as an Oxley Lagoon patch 2 plant falling close to the patch 1 samples at this site and Lake Noonamah previously mentioned. The third and fourth PCoA axes showed further clustering and highlighted further divergence of some patches. The third axis accounted for a further 10.6% of the total genetic variation and highlighted strong differentiation of samples collected in Lake Nooran patch 1, while the fourth axis highlighted strong differentiation of samples collected from Lake Noonamah patch 1 and accounted for a further 7.8% of the total variation (Figure 3b).

**Figure 3.**
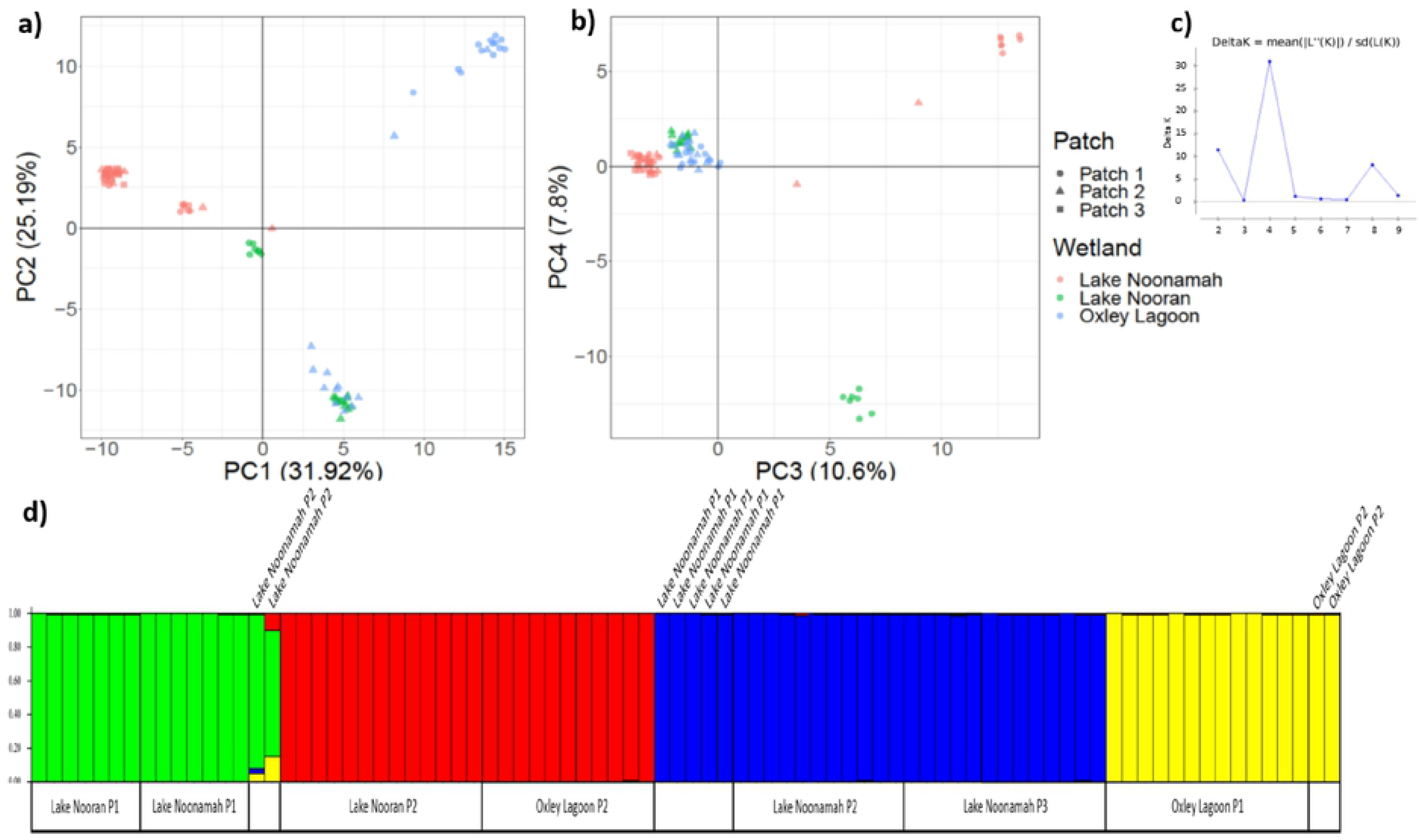
*a) Plot showing axis 1 and 2 and b) axis 3 and 4 from PCoA analysis of 84* Marsilea drummondii *genotypes across three wetlands (2-3 patches per wetland) in the lower Lachlan River Catchment, c) Representation of the Akaike value used to determine the number of populations (K), and d) Representation of the probability of population assignment for each individual genotype coloured by cluster and ordered by wetland and patch.*

Structure analysis showed four likely genetic clusters in *M. drummondii* (Figure 3c). The first cluster consisted of all individuals from Lake Nooran patch 1, seven samples from Lake Noonamah patch 1 and two samples from Lake Noonamah patch 2 (Figure 3d). The second cluster consisted of all samples from Lake Nooran patch 2 and all samples from Oxley Lagoon Patch 2. The third cluster consisted of four samples from Lake Noonamah patch 1, 11 (of 13) from Lake Noonamah patch 2, and all (13) samples from Lake Noonamah patch 3. The fourth cluster consisted of all (13) samples from Oxley Lagoon patch 1 and two samples from Oxley Lagoon patch 2.

The *E. acuta* PCoA axis 1 accounted for the majority of the variation (59.5%) and highlighted strong differentiation between samples collected from Lake Noonamah and those from Oxley Lagoon and Lake Nooran (Figure 4a). PCoA axis 2 accounted for 5% of the variation and highlighted differentiation between samples collected from Lake Nooran patch 1 and patch 2. The third axis (not shown) accounted for 4.2% of the variation and differentiation between the samples collected from Oxley Lagoon and Lake Nooran. This PCoA also highlighted strong within-patch clustering of samples collected from almost all of the patches. The structure analysis confirmed these two genetic clusters (Figure 4b), with the largest group consisting of all samples collected from Oxley Lagoon and Lake Nooran and the other group consisting of all 39 samples collected from within Lake Noonamah (Figure 4c).

**Figure 4.**
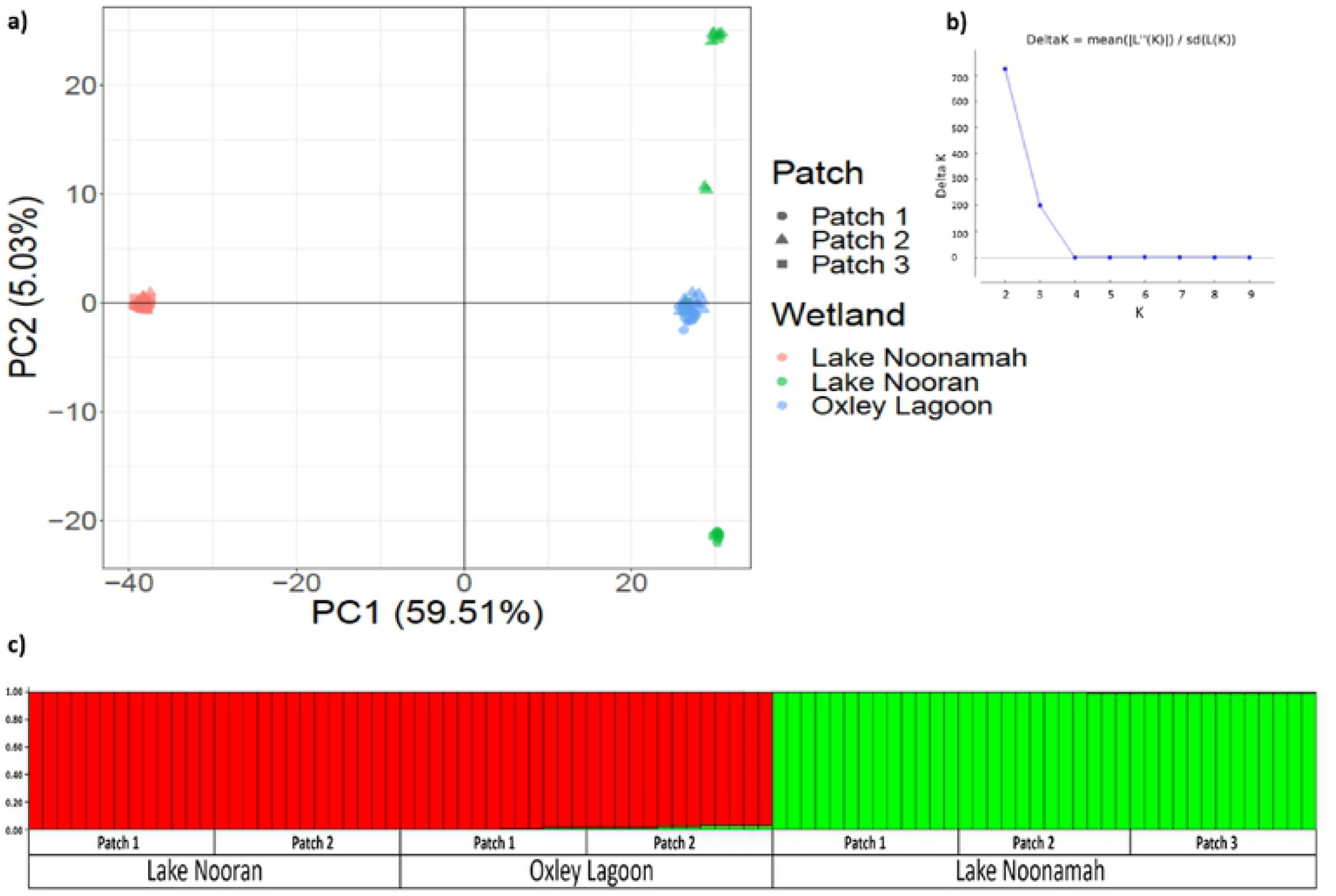
*a) Plot from PCoA analysis of 86* Eleocharis acuta *genotypes across three wetlands (2-3 patches per wetland) in the lower Lachlan River Catchment, b) Representation of the Akaike value used to determine the number of populations (K), and c) Representation of the probability of population assignment with each individual genotype coloured by cluster and ordered by wetland and patch.*

Hierarchical AMOVA demonstrated that the majority of variation was attributed to that within individuals (64% for *M. drummondii* and 67% for *E. acuta*) which was significant in both species (Table 4). A total of 31% and 32% of the variation in *M. drummondii* and *E. acuta* was attributed to variation among wetlands respectively, which was significant in both species (Table 4). Variation between samples within wetlands was estimated to be 5% in *M. drummondii* and 1% in *E. acuta* which were both non-significant (Table 4).

**Table 4.** *Analysis of molecular variance (AMOVA) for* Marsilea drummondii *and* Eleocharis acuta. *\* = significant p <0.05.*

| Source of variation | df | Sum of squares | Mean sum of squares | Estimated (%) variation | P value |
| --- | --- | --- | --- | --- | --- |
| *Marsilea drummondii* |  |  |  |  |  |
| Between wetlands | 2 | 7,092 | 3,546 | 31% | 0.01* |
| Between samples within wetlands | 81 | 12,227 | 150 | 5% | 0.12 |
| Within samples | 84 | 10,943 | 130 | 64% | 0.01* |
| Total | 167 | 30,262 | 181 | 100% |  |
| *Eleocharis acuta* |  |  |  |  |  |
| Between wetlands | 2 | 64,774 | 32,387 | 32% | 0.01* |
| Between samples within wetlands | 87 | 99,304 | 1,141 | 1% | 0.34 |
| Within samples | 90 | 98,384 | 1,093 | 67% | 0.01* |
| Total | 179 | 262,462 | 1,466 | 100% |  |

Pairwise *F*_ST_ values for *M. drummondii* ranged from 0.10 between Oxley Lagoon and Lake Nooran to 0.12 between Oxley Lagoon and Lake Noonamah (Table 5). Similarly, pairwise *F*_ST_ values in *E. acuta* ranged from 0.03 between Oxley Lagoon and Lake Nooran to 0.25 between Lake Nooran and Lake Noonamah. Pairwise *F*_ST_ between patches were higher in both species than comparisons at the wetland scale. For *M. drummondii, F*_ST_ ranged from almost no variation (*F*_ST_ = 0.03) between Lake Noonamah patch 2 and Lake Noonamah patch 3 to the highest difference between Lake Nooran patch 1 and Lake Noonamah patch 3 (*F*_ST_ = 0.40). For *E. acuta, F*_ST_ ranged from 0.03 between Lake Noonamah patches 1 and 2 to ∽0.31 between all Lake Noonamah patches and Lake Nooran patches 1 and 2 (Table 6).

**Table 5.** *Pairwise* F_*ST*_ *values for each wetland for* Marsilea drummondii *on the bottom right and* Eleocharis acuta *shaded in grey on the top left of the table.*

|  | Lake Nooran | Lake Noonamah | Oxley Lagoon |
| --- | --- | --- | --- |
| Lake Nooran |  | 0.25 | 0.03 |
| Lake Noonamah | 0.11 |  | 0.23 |
| Oxley Lagoon | 0.10 | 0.12 |  |

**Table 6.**
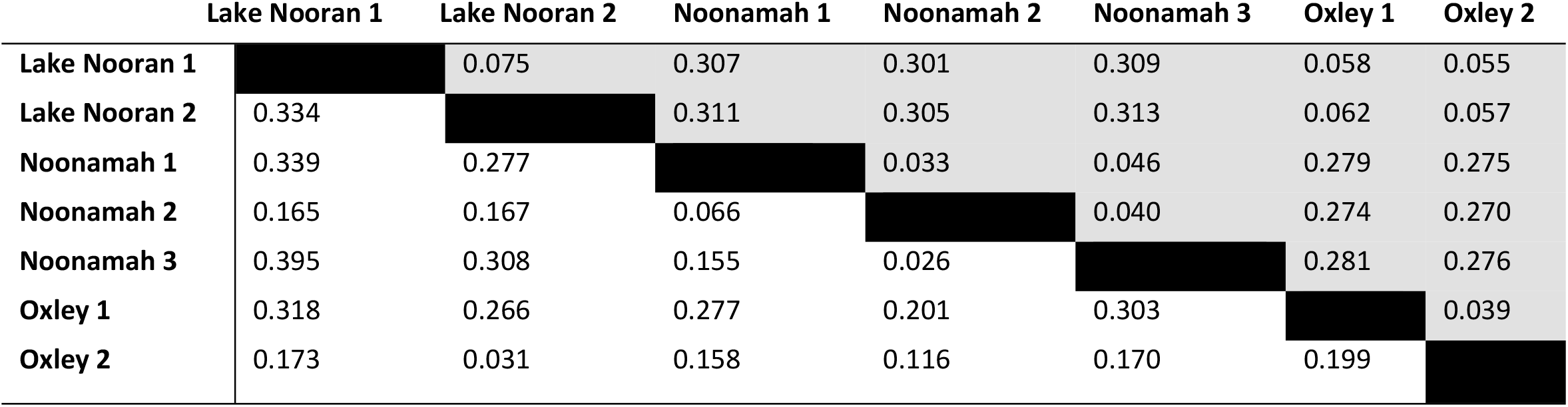
F_*ST*_ *values for each patch for* Marsilea drummondii *on the bottom right and* Eleocharis acuta *shaded in grey on the top left of the table*.

## Discussion

The results here demonstrate that, like most aquatic plants (Philbrick and Les 1996) asexual vegetative reproduction occurs in *M. drummondii* and *E. acuta* with multiple samples in both species having identical genotypes. Unexpectedly, however, the relatively low occurrence of identical genotypes in both species suggesting that this is not the major reproductive strategy of either species. Whilst *M. drummondii* and *E. acuta* grow clonally through rhizomes, the extent of clonality and size and arrangement of genotypes varied between the two species.

Seven samples contributed to the same genotype in *M. drummondii* which were collected in one patch within Lake Nooran, highlighting the potential extent (approximately 4 square metres) of a single genet (clone) in this species. This same genotype was also sampled from other patches within the same wetland highlighting the potential ability (albeit uncommon based on our data) of asexual dispersal via vegetative propagules within a wetland. In *E. acuta*, only one pair of identical samples were detected within a patch at a distance of 50 cm, demonstrating that vegetative reproduction also occurs in this species, however, the size of an individual genet is no more than 50 cm and likely to be most commonly smaller.

Although the results show that asexual vegetative reproduction does occur, dispersal of asexual propagules away from the parent plant is rare in *M. drummondii* and unlikely in *E. acuta*, and may be considered lower than what would be expected in aquatic macrophytes (Boedeltje *et al*. 2004; Li 2014). Boedeltje *et al*. (2004) found that 90% hydrochorous propagules in streams in the Netherlands were vegetative. However, many species showed a preference for either vegetative or generative dispersal, for example *Phragmites australis* was recorded predominately from generative (sexual) diaspores while *Potamogeton natans* exclusively by vegetative propagules (Boedeltje *et al*. 2008).

Vegetative growth in *E. acuta* and *M. drummondii* is through the production of rhizomes which have low dispersibility compared with other vegetative organs such as stolons, runners and bulbils (Barrett 2015). However, rhizomes of some aquatic plants, such as *Phragmites australis*, can float for at least six months under experimental conditions once removed from the soil (Sarneel 2013). For vegetative dispersal to occur in *E. acuta* and *M. drummondii* a combination of destructive removal (such as by birds) coinciding with flooding conditions may be required.

Sexual reproduction appears to be the dominant reproductive strategy in both species, and patches consist of numerous genets. This suggest that patches are regularly invaded by additional genotypes of sexual origin and that genets are rarely large and dominant. Both *E. acuta* and *M. drummondii* have persistent soil seed or spore banks (Lellinger 1985; Higgisson and Dyer 2021) and these results suggest a reliance on these soil banks rather than on vegetative reproduction for population persistence.

The reliance on sexual reproduction rather than vegetative shown here may be related to the infrequent and variable flooding regime, and extended dry-phases experienced on river-floodplain systems in dryland regions such as the lower Lachlan River (Higgisson *et al*. 2020), compared with aquatic plants in more permanent hydrological regimes. Persistent soil seedbanks are known to be important in enabling aquatic plants to persevere on floodplains in dryland regions (Capon and Brock 2006). A similar situation is likely to exist in the water fern examined here since *M. strigosa* growing in the Mediterranean basin was found to rely on sexual rather than vegetative reproduction, with the authors suggesting that sporocarps were the only means for this species to survive the harsh winter conditions (Vitalis *et al*. 2002). The results from this study demonstrate the likely importance of sexual reproduction and the ability of sporocarps and seeds to remain viable during often extended dry periods for the survival of *M. drummondii* and *E. acuta* in dryland river floodplains. The longevity of *E. acuta* seeds and *M. drummondii* sporocarps may help to reduce genetic drift and retain genetic variation in populations.

Like self-compatible flowering plants, heterosporous ferns such as *M. drummondii* produce both male and female sex organs, megaspores (male spores) and microspores (female spores), and as such are partially hermaphroditic and have the ability to self-fertilize (Schneider and Pryer 2002). As these are heterosporous, intragametophytic self-fertilisation (sperm and egg from one gametophyte producing the sporophyte) is not possible since the egg and sperm are on separate gametophytes.

However, intergametophytic self-fertilisation involving separate gametophytes derived from the one sporophyte can occur. Our relatedness analyses provides strong evidence that *M. drummondii* can and does reproduce via self-fertilisation with nine samples primarily from the same site in parent-offspring relationships via self-fertilisation. Self-fertilisation has been demonstrated in *M. strigosa* in the Mediterranean (Vitalis *et al*. 2002). Interestingly, reproduction via self-fertilisation only occurred within Lake Noonamah highlighting that differences between wetlands in landscape and hydrological processes may influence the rate of self-fertilisation in *M. drummondii* and potentially other species where reproduction occurs in water. Among population variation in plant mating systems (self-fertilisation vs exclusive outcrossing) within species has been observed in other plants (Whitehead *et al*. 2018).

In *E. acuta*, only one pair of samples were in a parent-offspring relationship through self-fertilisation, suggesting that while this species can reproduce via self-fertilisation, mechanisms to reduce self-fertilisation such as dichogamy, observed in *E. laeviglumis* (Demeda *et al*. 2018) exist. Other mechanisms to promote outcrossing also exist in the Cyperaceae such as self-incompatibility in *Rhynchospora ciliata* (Costa and Machado 2012).

During a reproductive experiment of multiple *Marsilea* species, Schneider and Pryer (2002) observed that submerged sporocarps sank then opened within a few hours followed by the simultaneous release of megaspores and microspores which drifted to the water surface where fertilisation occurred over approximately 10 hours. This suggests that *M. drummondii* has no apparent adaptation to reduce selfing, apart from the opportunistic mixing of spores. The greater the number of spores from different genotypes on the water surface the more likely is cross-pollination.

The measures of within-wetland genetic diversity found in *M. drummondii* and *E. acuta* were both highest in Oxley Lagoon, intermediate in Lake Nooran and lowest in Lake Noonamah. These similar patterns despite different life-histories, highlight how genetic diversity in aquatic plants in dryland river floodplains are driven by numerous factors including geographic barriers and hydrological regime. Oxley Lagoon is situated in very close proximity to the Lachlan River (<50 m), Lake Nooran is approx. 200 m away, while Lake Noonamah is >5km from the main channel suggesting that distance from the river may influence levels of genetic diversity. Genetic diversity was found to increase in down-stream populations of *Sparganium emersum* (unbranched bur-reed), related to the unidirectional downstream flow on diversity patterns (Pollux *et al*. 2009). Similarly, our results suggest that genetic diversity decreases the further from the main channel a population is located.

The high levels of genetic differentiation and clustering within patches and wetlands suggests that pollination and dispersal are most likely in close proximity to parent plants. Colonisation (founder) events between wetlands appear to be very rare in *E. acuta* but more frequent in *M. drummondii*. This high genetic differentiation further suggests that colonisation (or recolonisation following localised extinction) occurs from the local propagule pool (deme/wetland) and mixing of colonising individuals from different demes is unlikely (Wade 1978). The limited geneflow from either sexual or asexual propagules in these semi-aquatic species is likely a key driver in population genetic structure by promoting genetic differentiation. The possible influence of local conditions in the fixing of alleles in different wetlands and even different patches (following Slatkin 1987) as well as founder effects are also likely to infleunce the population genetic structure observed.

Fifty percent of *E. acuta* seeds floated for at least 30 days, in an experiment by Higgisson and Dyer (2021), indicating a capacity for long distance dispersal under suitable conditions. Whilst potential dispersal maybe much greater in *E. acuta*, here we show that realized seed dispersal is most likely within proximity to the parent plant and within a wetland. Although hydrochorous dispersal of sporocarps is unknown, these start to open within a few hours of contact with water (Schneider and Pryer 2002) possibly limiting long distance hydrochory. While our results suggest that dispersal is most likely to occur within proximity of the parent plant, the finding that some *M. drummondii* individuals were scattered beyond the patch they were sampled in the PCoA, suggests gene flow is occurring between wetlands; this situation was not observed in *E. acuta*.

The floodplain of the lower Lachlan River is known as a nesting and feeding habitat for a range of waterbirds, which have been recorded traveling along the Lachlan River and between its wetlands (McGinness 2021). In the absence of specific information on *M. drummondii*, the sporocarps of *M. mucronata* have been shown to pass intact through the digestive tract of waterbirds (Malone and Proctor 1965) showing the likely role of waterbirds in their dispersal. Whilst intact seeds of *E. acuta* were recorded in fecal samples of the waterbird Grey Teal, none of these seeds germinated during experimental trials (Green *et al*. 2008). Lowered seed viability following ingestion be waterbirds may have contributed to the lower connectivity and gene flow between wetlands shown here in *E. acuta* compared with *M. drummondii*.

### Implications

While asexual vegetative reproduction does occur in *M. drummondii* and *E. acuta*, sexual reproduction is the dominant reproductive strategy. Dispersal between wetlands (and even between patches) is infrequent in *M. drummondii* and extremely rare in *E. acuta*. This suggests a reliance on local seed sources and seed or spore bank longevity. Low dispersal means recolonizing wetland patches is challenging especially under a drying climate across the study region where the timing, frequency, and volume of precipitation is likely to dramatically change. In river-floodplain systems in dryland regions such as the lower Lachlan River, the recolonisation of patches following disturbance such as an extended drought maybe unlikely. These results highlight the importance of protecting existing populations and maintaining diverse and functioning wetlands. There may also be a need to collect and store propagules for future restoration activities. Once lost from a wetland these species may not have the ability to recolonize, especially under regulated flow conditions.

## Acknowledgements

The authors would like to thank Yasmin Cross, Tasha James, Jasmin Wells and Matt Young for assisting with the collection of plant samples and Alica Tschierschke for providing the map of the study area. The authors also thank the landholders who provided us access to their properties to collect the leaf samples. This research was funded by the Centre for Applied Water Science, University of Canberra.

## Data Accessibility

The genetic and meta data have been provided to the journal as part of the submission of this manuscript and has also been archived in the University of Canberra, Institute for Applied Ecology data repository and is available upon request. The R code which we developed to determine the pairwise samples in each of the three relatedness scenarios is also available upon request.

## Author Contributions

- William Higgisson – conceived and designed the study, undertook the field work, analyzed the data and wrote the paper
- Linda Broadhurst – wrote the paper
- Foyez Shams – analyzed the data
- Burnd Gruber – analyzed the data
- Fiona Dyer – Designed the study and wrote the paper

## References

Altshuler, D., Pollara, V.J., Cowles, C.R., Van Etten, W.J., Baldwin, J., Linton, L., and Lander, E.S. (2000). An SNP map of the human genome generated by reduced representation shotgun sequencing. Nature, 407, 513–516.

Baird, N.A., Etter, P.D., Atwood, T.S., Currey, M.C., Shiver, A.L., Lewis, Z.A., Selker, E.U., Cresko, W.A., and Johnson, E.A. (2008). Rapid SNP discovery and genetic mapping using sequenced RAD markers. PloS one, 3, e3376.

Barrett, S.C. (1980). Sexual reproduction in Eichhornia crassipes (water hyacinth). II. Seed production in natural populations. Journal of applied ecology, 113–124.

Barrett, S.C. (2015). Influences of clonality on plant sexual reproduction. Proceedings of the National Academy of Sciences, 112, 8859–8866.

Boedeltje, G., Bakker, J.P., Ten Brinke, A., Van Groenendael, J.M., and Soesbergen, M. (2004). Dispersal phenology of hydrochorous plants in relation to discharge, seed release time and buoyancy of seeds: the flood pulse concept supported. Journal of Ecology, 92, 786–796.

Boedeltje, G., Ozinga, W.A., and Prinzing, A. (2008). The trade-off between vegetative and generative reproduction among angiosperms influences regional hydrochorous propagule pressure. Global Ecology and Biogeography, 17, 50–58.

Bureau of Meteorology (2021a). ‘Climate statistics for Australian locations, Summary statistics Oxley (Walmer Downs)’. Available at http://www.bom.gov.au/climate/data-services/station-data.shtml [accessed 21/06/2021].

Bureau of Meteorology (2021b). ‘Monthly mean maximum temperature, Hay Airport AWS’. Available at [accessed 21/06/2021].

Cain, M.L., Milligan, B.G., and Strand, A.E. (2000). Long-distance seed dispersal in plant populations. American journal of botany, 87, 1217–1227.

Capon, S.J., and Brock, M.A. (2006). Flooding, soil seed bank dynamics and vegetation resilience of a hydrologically variable desert floodplain. Freshwater Biology, 51, 206–223.

Costa, A., and Machado, I. (2012). Flowering dynamics and pollination system of the sedge Rhynchospora ciliata (Vahl) Kükenth (Cyperaceae): does ambophily enhance its reproductive success? Plant Biology, 14, 881–887.

Cruz, V.M., Kilian, A., and Dierig, D.A. (2013). Development of DArT marker platforms and genetic diversity assessment of the US collection of the new oilseed crop lesquerella and related species. PLoS One, 8, e64062.

Cunningham, G., Mulham, W., Milthorpe, P., and Leigh, J. (1981). ‘Plants of Western New South Wales.’ (CSIRO Publishing: Melbourne, Vic., Australia.)

Demeda, C.L.B., Seger, G.D.d.S., Steiner, N., and Trevisan, R. (2018). Reproductive phenology and germination of Eleocharis laeviglumis R. Trevis. & Boldrini (Cyperaceae). Acta Botanica Brasilica, 32, 487–492.

Dorken, M.E., and Eckert, C.G. (2001). Severely reduced sexual reproduction in northern populations of a clonal plant, Decodon verticillatus (Lythraceae). Journal of Ecology, 339–350.

Earl, D.A. (2012). STRUCTURE HARVESTER: a website and program for visualizing STRUCTURE output and implementing the Evanno method. Conservation genetics resources, 4, 359–361.

Eckert, C.G., Dorken, M.E., and Mitchell, S.A. (1999). Loss of sex in clonal populations of a flowering plant, Decodon verticillatus (Lythraceae). Evolution, 53, 1079–1092.

Eckert, C.G., Dorken, M.E., and Barrett, S.C. (2016). Ecological and evolutionary consequences of sexual and clonal reproduction in aquatic plants. Aquatic Botany, 135, 46–61.

Elshire, R.J., Glaubitz, J.C., Sun, Q., Poland, J.A., Kawamoto, K., Buckler, E.S., and Mitchell, S.E. (2011). A robust, simple genotyping-by-sequencing (GBS) approach for high diversity species. PloS one, 6, e19379.

Evanno, G., Regnaut, S., and Goudet, J. (2005). Detecting the number of clusters of individuals using the software STRUCTURE: a simulation study. Molecular Ecology, 14, 2611–2620.

Goudet, J. (2005). Hierfstat, a package for R to compute and test hierarchical F-statistics. Molecular Ecology Notes, 5, 184–186.

Green, A.J., Jenkins, K., Bell, D., Morris, P., and Kingsford, R. (2008). The potential role of waterbirds in dispersing invertebrates and plants in arid Australia. Freshwater Biology, 53, 380–392.

Gruber, B., Unmack, P.J., Berry, O.F., and Georges, A. (2018). dartr: An r package to facilitate analysis of SNP data generated from reduced representation genome sequencing. Molecular Ecology Resources, 18, 691–699.

Handel, S.N. (1985). The intrusion of clonal growth patterns on plant breeding systems. The American Naturalist, 125, 367–384.

Hanski, I. (1998). Metapopulation dynamics. Nature, 396, 41–49.

Harper, J.L. (1977). Population biology of plants. Population biology of plants.

Higgisson, W., Higgisson, B., Powell, M., Driver, P., and Dyer, F. (2020). Impacts of water resource development on hydrological connectivity of different floodplain habitats in a highly variable system. River research and applications, 36, 542–552.

Higgisson, W., and Dyer, F. (2021). Seed germination and dispersal of Eleocharis acuta and Eleocharis sphacelata under experimental hydrological conditions. Aquatic Ecology, 1–12.

Ivey, C.T., and Richards, J.H. (2001). Genetic diversity of everglades sawgrass, Cladium jamaicense (Cyperaceae). International Journal of Plant Sciences, 162, 817–825.

Johnson, D.M. (1986). Systematics of the new world species of Marsilea (Marsileaceae). Systematic Botany Monographs, 1-87.

Jones, D.L. (1998). Marsileaceae. In Flora of Australia: Ferns, gymnosperms and allied groups. pp. pp.166–173. (CSIRO Publishing: Australia.)

Junk, W.J., Bayley, P.B., and Sparks, R.E. (1989). The flood pulse concept in river-floodplain systems. Canadian Special Publication of Fisheries and Aquatic Sciences, 106, 110–127.

Kamvar, Z., Tabima, J., and Grünwald, N. (2014). Poppr: an R package for genetic analysis of populations with clonal, partially clonal, and/or sexual reproduction. PeerJ., 2: e281

Kilian, A., Wenzl, P., Huttner, E., Carling, J., Xia, L., Blois, H., Caig, V., Heller-Uszynska, K., Jaccoud, D., and Hopper, C. (2012). Diversity arrays technology: a generic genome profiling technology on open platforms. Methods in Molecular Biology, 888, 67–89.

Kingsford, R.T. (2000). Ecological impacts of dams, water diversions and river management on floodplain wetlands in Australia. Austral Ecology, 25, 109–127.

Le Corre, V., Machon, N., Petit, R.J., and Kremer, A. (1997). Colonization with long-distance seed dispersal and genetic structure of maternally inherited genes in forest trees: a simulation study. Genetics Research, 69, 117–125.

Lellinger, K. (1985). New records for longevity of Marsilea sporocarps. American Fern Journal, 75, 30–31.

Les, D.H. (1988). Breeding systems, population structure, and evolution in hydrophilous angiosperms. Annals of the Missouri Botanical Garden, 819–835.

Li, W. (2014). Environmental opportunities and constraints in the reproduction and dispersal of aquatic plants. Aquatic Botany, 118, 62–70.

Liu, L., Wang, J., Ma, X., Li, M., Guo, X., Yin, M., Cai, Y., Yu, X., Du, N., and Wang, R. (2021). Impacts of the yellow River and Qingtongxia dams on genetic diversity of Phragmites australis in Ningxia Plain, China. Aquatic Botany, 169, 103341.

Malone, C.R., and Proctor, V.W. (1965). Dispersal of Marsilea mucronata by water birds. American Fern Journal, 55, 167–170.

McGinness, H. (2021). ‘Waterbird breeding and movements: Knowledge for water managers’. Available at https://research.csiro.au/ewkrwaterbirds/research-results/ [accessed 5/05/2021].

McKee, J., and Richards, A. (1996). Variation in seed production and germinability in common reed (Phragmites australis) in Britain and France with respect to climate. New Phytologist, 133, 233–243.

MDBA (2012). ‘Assessment of environmental water requirements for the proposed Basin Plan. Series of 24 reports, Australian Government, Murray-Darling Basin Authority’. Available at https://www.mdba.gov.au/publications/mdba-reports/assessing-environmental-water-requirements-basins-rivers [accessed 6 May 2022].

Nagalingum, N.S., Schneider, H., and Pryer, K.M. (2006). Comparative morphology of reproductive structures in heterosporous water ferns and a reevaluation of the sporocarp. International Journal of Plant Sciences, 167, 805–815.

Niklas, K.J., and Cobb, E.D. (2017). The evolutionary ecology (evo-eco) of plant asexual reproduction. Evolutionary Ecology, 31, 317–332.

Ozinga, W.A., Bekker, R.M., Schaminee, J.H., and Van Groenendael, J.M. (2004). Dispersal potential in plant communities depends on environmental conditions. Journal of Ecology, 92, 767–777.

Pannell, J.R., and Charlesworth, B. (1999). Neutral genetic diversity in a metapopulation with recurrent local extinction and recolonization. Evolution, 53, 664–676.

Peakall, R., and Smouse, P.E. (2006). GENALEX 6: genetic analysis in Excel. Population genetic software for teaching and research. Molecular Ecology Notes, 6, 288–295.

Philbrick, C.T., and Les, D.H. (1996). Evolution of aquatic angiosperm reproductive systems. BioScience, 46, 813–826.

Piquot, Y., Saumitou-Laprade, P., Petit, D., Vernet, P., and Epplen, J. (1996). Genotypic diversity revealed by allozymes and oligonucleotide DNA fingerprinting in French populations of the aquatic macrophyte Sparganium erectum. Molecular Ecology, 5, 251–258.

PlantNET (2021a). ‘The NSW Plant Information Network System, Royal Botanic Gardens and Domain Trust, Sydney ‘. Available at [accessed 13/02/2020].

PlantNET (2021b). ‘The NSW Plant Information Network System’. Available at [accessed 06/01/2021].

Pollux, B., Luteijn, A., Van Groenendael, J., and Ouborg, N. (2009). Gene flow and genetic structure of the aquatic macrophyte Sparganium emersum in a linear unidirectional river. Freshwater Biology, 54, 64–76.

Pritchard, J.K., Stephens, M., and Donnelly, P. (2000). Inference of population structure using multilocus genotype data. Genetics, 155, 945–959.

Reynes, L., Thibaut, T., Mauger, S., Blanfuné, A., Holon, F., Cruaud, C., Couloux, A., Valero, M., and Aurelle, D. (2020). Genomic signatures of clonality in the deep water kelp Laminaria rodriguezii. Authorea Preprints

Santamaría, L. (2002). Why are most aquatic plants widely distributed? Dispersal, clonal growth and small-scale heterogeneity in a stressful environment. Acta oecologica, 23, 137–154.

Sarneel, J. (2013). The dispersal capacity of vegetative propagules of riparian fen species. Hydrobiologia, 710, 219–225.

Schaible, R., Bergmann, I., Bögle, M., Schoor, A., and Schubert, H. (2009). Genetic characterisation of sexually and parthenogenetically reproductive populations of Chara canescens (Charophyceae) using AFLP, rbc L, and SNP markers. Phycologia, 48, 105–117.

Schneider, H., and Pryer, K.M. (2002). Structure and function of spores in the aquatic heterosporous fern family Marsileaceae. International Journal of Plant Sciences, 163, 485–505.

Slatkin, M. (1987). Gene flow and the geographic structure of natural populations. Science, 236, 787–792.

Van der Valk, A. (1981). Succession in wetlands: a Gleasonian approach. Ecology, 688-696.

Vitalis, R., Riba, M., Colas, B., Grillas, P., and Olivieri, I. (2002). Multilocus genetic structure at contrasted spatial scales of the endangered water fern Marsilea strigosa Willd.(Marsileaceae, Pteridophyta). American journal of botany, 89, 1142–1155.

Wade, M.J. (1978). A critical review of the models of group selection. The Quarterly Review of Biology, 53, 101–114.

Walker, K.F., Sheldon, F., and Puckridge, J.T. (1995). A perspective on dryland river ecosystems. Regulated Rivers: Research & Management, 11, 85–104.

Walters, S. (1949). Eleocharis R. Br. The Journal of Ecology, 192–206.

Whitehead, M.R., Lanfear, R., Mitchell, R.J., and Karron, J.D. (2018). Plant mating systems often vary widely among populations. Frontiers in Ecology and Evolution, 6, 38.

